# Long-read RNA sequencing of human and animal filarial parasites improves gene models and discovers operons

**DOI:** 10.1101/2020.06.18.160267

**Authors:** Nicolas J Wheeler, Paul M. Airs, Mostafa Zamanian

## Abstract

Filarial nematodes (Filarioidea) cause substantial disease burden to humans and animals around the world. Recently there has been a coordinated global effort to generate and curate genomic data from nematode species of medical and veterinary importance. This has resulted in two chromosome-level assemblies (*Brugia malayi* and *Onchocerca volvulus*) and 10 additional draft genomes from Filarioidea. These reference assemblies facilitate comparative genomics to explore basic helminth biology and prioritize new drug and vaccine targets. While the continual improvement of genome contiguity and completeness advances these goals, experimental functional annotation of genes is often hindered by poor gene models. Short-read RNA sequencing data and expressed sequence tags, in cooperation with *ab initio* prediction algorithms, are employed for gene prediction, but these can result in missing clade-specific genes, fragmented models, imperfect mapping of gene ends, and lack of isoform resolution. Long-read RNA sequencing can overcome these drawbacks and greatly improve gene model quality. Here, we present Iso-Seq data for *B. malayi* and *Dirofilaria immitis*, etiological agents of lymphatic filariasis and canine heartworm disease, respectively. These data cover approximately half of the known coding genomes and substantially improve gene models by extending untranslated regions, cataloging novel splice junctions from novel isoforms, and correcting mispredicted junctions. Furthermore, we validated computationally predicted operons, identified new operons, and merged fragmented gene models. We carried out analyses of poly(A) tails in both species, leading to the identification of non-canonical poly(A) signals. Finally, we prioritized and assessed known and putative anthelmintic targets, correcting or validating gene models for molecular cloning and target-based antiparasitic screening efforts. Overall, these data significantly improve the catalog of gene models for two important parasites, and they demonstrate how long-read RNA sequencing should be prioritized for future improvement of parasitic nematode genome assemblies.

## Introduction

Infections caused by parasitic filarial nematodes inflict massive burdens upon humans and companion animals around the world. The mosquito-borne filarial worm *Brugia malayi* is an etiological agent of the neglected tropical disease lymphatic filariasis (LF), which affects over 60 million people worldwide [1]. *Dirofilaria immitis*, a related mosquito-borne filarial worm, is a causative agent of dog heartworm and a zoonotic concern [2]. These diseases are ultimately controlled by different strategies of preventive chemotherapy. In tropical regions endemic for LF, the World Health Organization has recommended mass drug administration (MDA) of two annual doses of albendazole (ABZ) or a combination of ABZ, ivermectin (IVM), and/or diethylcarbamazine citrate (DEC) [3]. For control of dog heartworm, the American Heartworm Society recommends monthly administration of macrocyclic lactones like IVM [4].

Preventive chemotherapies are active against parasite larvae; no adulticidal therapies are approved for the treatment of LF, and only the arsenic-containing drug melarsomine is available for clearing adult heartworm. Further, while the triple-drug combination is a new development for LF control, the absence of novel filaricidal chemical moieties combined with the massive drug pressure caused by preventive chemotherapy at a global scale continues to increase fear of loss of drug efficacy for both the human and veterinary diseases [5–7]. Indeed, this is already an emerging problem for heartworm control [8,9].

LF and heartworm control and elimination strategies would be significantly helped with the development of new antifilarial drugs and therapies. New data and technologies have promised to speed the discovery and validation of new therapeutic targets. The release of the *B. malayi* draft genome initiated a wave of target-based approaches that leverage genomic data, molecular biology, and rational target selection, which allow for the cloning, expression, and screening of putative targets in recombinant systems or exploring the repurposing of approved drugs with known targets [10–13]. Intrinsic to this process is the assumption that the desired transcript and isoform of a given target is known and amenable to straightforward cloning.

However, while the completeness of the *B. malayi* and *D. immitis* draft genomes is sufficient for target prioritization and selection via comparative genomic approaches, the fragmentation and incompleteness of predicted gene models can delay facile cloning, expression, and characterization or screening (see [10], for instance). Despite the excellent chromosome-level assembly of the *B. malayi* genome [14], two of three cloned genes in a recent study [15] had incorrectly predicted splice junctions even though these genes share high similarity with their one-to-one *C. elegans* homologs. Gene prediction and curation in *Caenorhabditis* spp. is known to be a difficult task, and it is clear that is also the case in filarial nematodes [16].

In addition to incorrect gene models, both *B. malayi* and *D. immitis* have very few genes with predicted isoforms. *C. elegans* (WS277) currently has 1.97 predicted transcripts per gene, while *B. malayi* (WS277) has 1.44 and *D. immitis* (WBPS14) does not have a single predicted isoform. It is increasingly clear that many anthelmintics have selectivity for specific target isoforms. IVM exhibits strong isoform-specific action against *C. elegans* AVR-14 ion channel subunits expressed in *Xenopus* [17], and this has been validated in a range of parasitic nematodes, including *D. immitis* [18–20]. More recently, a target of emodepside, the calcium channel SLO-1, was shown to cause sex-specific effects in adult *B. malayi* due to alternative splicing of the target [21]. Thus, identifying transcript isoforms should be a high priority in the continual curation of the genomes of parasitic nematodes.

A number of techniques are available to generate datasets that aid supervised gene prediction algorithms. Cap analysis gene expression (CAGE) has been used to characterize the landscape of transcriptional start sites (TSS) in a given species or tissue [22]. However, many nematode pre-mRNAs lose their original 5’ end through *trans*-splicing to splice-leader (SL) sequences (∼70% in *B. malayi* [23]), obscuring the true TSS and making the CAGE technique unsuitable. In *C. elegans*, a modified technique has been used that leveraged the ability to rear worms at lower temperatures to slow the processing of *trans*-spliced transcripts [24], but most parasitic nematodes are not amenable to these modifications in animal husbandry. Additionally, the integration of ever-increasing short-read RNA sequencing data provides support for exon usage, but it is difficult to generate abundant high-quality mRNA across many parasitic nematode life stages. This difficulty has resulted in databases with a strong bias in reads mapping to the 3’ end of transcripts. Even where RNA quality and yield are not limiting, alternative splicing and isoform prediction are not straightforward with exclusively short-read datasets.

Only two long-read RNA sequencing datasets exist for parasitic nematodes, the zoonotic hookworm *Ancylostoma ceylanicum* and the barber’s pole worm *Haemonchus contortus* [25,26]. Here, we report long-read RNA sequencing of the filarial parasites *B. malayi* and *D. immitis*. We separately sequenced both sexes of these species, allowing for the analysis of sex-specific isoform usage. We show a substantial increase in the lengths of predicted untranslated regions, identify operons, correct gene models, and catalog the parameters of poly(A) tail usage. This resource will significantly improve the confidence in gene predictions in these organisms and enhance the utility of the reference genomes.

## Methods

### Protocol and data availability

All data analysis and visualization pipelines are publicly available at https://github.com/zamanianlab/FilaridIsoSeq-ms. Long-read sequencing data has been deposited into NIH BioProjects PRJNA548902 (*B. malayi*) and PRJNA640410 (*D. immitis*).

### Parasite source

*B. malayi* adult males and females were sent to the University of Wisconsin-Madison from the Filariasis Reagent Resource Center (FR3)[27], washed, and incubated at 37°C in fresh complete media (RPMI 1640, 10% FBS, gentamicin, and penicillin/streptomycin) overnight prior to RNA extraction. *D. immitis* adult males and females were received at the University of Wisconsin-Madison from Zoetis, washed, and incubated at 39°C in fresh complete media (RPMI 1640, 20 mM HEPES, 1% D-glucose, 1% HCO_3_, 10% FBS, and penicillin/streptomycin). After overnight incubation, worms were individually dipped in PBS, transferred to an empty 50 mL conical tube, and flash-frozen in liquid N_2_. Frozen worms were sent to Smith College for RNA extraction.

### RNA extraction

15 *B. malayi* worms from each sex were combined and transferred to 750 uL TRIzol LS with 250 uL nuclease-free water. Samples were incubated for 1 hour prior to homogenization in a TissueLyser II for two cycles of 3 minutes at 30 Hz separated by a 2 minute incubation on ice. RNA was extracted according to the manufacturer’s recommendation, DNase treated, and re-purified with phenol:chloroform. RNA pellets were resuspended in nuclease-free water and assessed for quality and concentration with a Bioanalyzer 2100 (Agilent, Santa Clara, CA) RNA Pico chip. RNA was extracted from a single *D. immitis* worm from each sex using the PureLink RNA Mini kit (Thermo Fisher Scientific, Waltham, MA). RNA was DNase treated and subsequently dialyzed and assessed for quality and concentration with a Bioanalyzer 2100 (Agilent) RNA Pico chip. Extraction using these protocols provided RNA with RNA integrity (RIN) values between 6 and 8.

### SMRTbell library preparation, quality control, and sequencing

cDNA was synthesized from parasite RNA using the SMARTer PCR cDNA synthesis kit (Takara, Mountain View, CA), which incorporates a poly(A) selection step. 1 µg of RNA input was used and replicate cDNA reactions were performed where excess RNA was available. PCR cycles were optimized, and 12 cycles were used for all 4 samples. PCR reactions were pooled at 1X and 0.5X for each library, cleaned with 1X and 0.5X AMPure XP beads, and quantified on a 2100 Bioanalyzer (Agilent) with an HS DNA chip. Equimolar fractions of 1X and 0.5X cDNA were pooled with a target of >2 µg of each fraction, and libraries were created with the SMARTbell Template Library kit (Pacific Biosciences, Menlo Park, CA). The final library quality and concentration were assessed using a Qubit DNA HS kit (Thermo Fisher Scientific) and a 2100 Bioanalyzer (Agilent) HS DNA chip. A single SMRT cell was used for each library and sequenced on the Sequel system (PacBio) with diffusion loading and a 20 hour movie.

### Sequencing quality control and transcriptome analysis

Circular consensus sequences (CCSs) were generated from subreads with IsoSeq v3.1.0 using the genome assemblies from WormBase ParaSite (WBPS14) [14,28,29]. Transcripts were filtered with SQANTI3, which reference-corrects all transcripts and filters them based on hallmarks of artifacts and SVM classifiers [30]. Sequencing summary statistics were generated with SAMtools [31], and read depth was calculated with BEDtools [32,33].

### Identification of novel genes

We filtered CCSs to maintain those categorized as antisense (aligning to the opposite strand of a genomic locus with a predicted gene) and intergenic (aligning to a genomic locus without a predicted gene). These CCSs were used as queries for blastx [34] searches against the respective predicted proteomes [14,29]. CCSs with blast hits with e-values > 1e-3 (i.e., no significant hits to a predicted protein) were retained and then used as queries for blastx searches against the predicted proteomes of *Brugia pahangi* [35], *Caenorhabditis elegans* [36], *Ascaris suum* [37], *Onchocerca volvulus* [38], *Strongyloides ratti* [39], *and Toxocara canis* [40]. CCSs with blast hits with e-values < 1e-3 (i.e., significant hits to predicted proteins from other nematodes) were retained and finally used as queries in a blastn search against the *B. malayi* genomic scaffolds to determine the number of unique loci mapped.

### Analysis of poly(A) tails

We developed a single script to analyze poly(A) tails in untrimmed CCSs. Analyses include (1) measurement of poly(A) tail length, (2) discovery and tally of k-mers in the region upstream of the 3’ cleavage site, (3) identification of the most likely poly(A) signal (PAS), (4) location of the PAS, and (5) generation of a position-specific score matrix (PSSM) to calculate nucleotide frequencies at every position in the region flanking the 3’ cleavage site. The tool is open-source and utilizes open-source Python libraries.

Poly(A) tails are measured by counting A nucleotides until reaching a non-A dinucleotide. To identify k-mers enriched in 3’ UTRs, k-mers from the translational stop site to the 3’ cleavage site were tallied and compared to k-mer frequencies across all non-coding intergenic sequences from the relevant genome; this comparison resulted in a list of k-mers enriched in 3’ UTRs. The most likely PAS was identified by iteratively searching the upstream region for the k-mers from this enriched list, sorted by frequency; the search was terminated when a k-mer was found and the location of the PAS relative to the cleavage site was recorded. Finally, to analyze nucleotide frequencies in the region immediately upstream of the cleavage site for a given PAS, the poly(A) tail was trimmed from each sequence, CCSs were pooled by PAS, and the 50 bp upstream of the cleavage site were aligned. The PSSM function of BioPython [41] was implemented to calculate nucleotide frequencies at each position of the multiple sequence alignment.

### Operon identification

The GTF file for *B. malayi* was downloaded from WormBase ParaSite (WBPS14) [28], converted to BED format with the gtf2bed script from BEDOPS v2.4.37 [42], and “gene” features were retained. CCSs were aligned to reference genomes with minimap2 [43] and BAM files were converted to BED format with bamtobed from BEDTOOLS with the -split flag [32,33]. BEDTOOLS “intersect” with the -wa and -wb flags were used to retain exonic regions to which CCSs had been mapped. BED files were imported into R for further analysis.

Any CCS that mapped to more than one gene was kept as a potential polycistronic CCS. The distance between cistrons was calculated by taking the difference between the reference 3’ end of the 5’ coding sequence (CDS) and the reference 5’ end of the 3’ CDS; thus, incorrect reference annotations of the gene ends inevitably caused some errors in the distance calculation. We kept all putative operons that had a maximum intercistronic distance of <5000 nt, as the vast majority of assembly-annotated operons had intergenic distances below this threshold. We manually inspected each putative operon and made true/false decisions using a variety of evidence, including 1) the number of ORFs in CCSs spanning the genes (calculated using the ExPASy translate server [44]), 2) the sequence similarity of ORFs within CCSs to nematode orthologs (calculated using blastx on WormBase [45] or WormBase ParaSite [28]), and the 3) differences in lengths between these orthologs.

### Analysis of operon expression

Short-read RNA sequencing data from across the *B. malayi* life cycle [46] were acquired from NCBI SRA; reads were remapped to Bmal-4.0 and counts were generated with HISAT2 [47] and StringTie2 [48]. The RNA-seq pipeline was implemented using Nextflow [49] and is publicly available (https://github.com/zamanianlab/BmalayiRNAseq-nf).

Only the first two cistrons of each operon were kept, and pseudo-operons were created by randomly selecting neighboring genes with intergenic distances <5000 nt. Expression abundances for each gene pair - putative polycistrons and pseudo-operons - during each life cycle stage (22 RNA-seq data sets) were fit using a linear model, and R^2^ values were calculated.

### Drug target analysis

We generated a curated list of genes that are known anthelmintic targets and receptors belonging to druggable protein families in parasitic nematodes, including selected ligand-gated ion channels (LGICs), voltage-gated ion channels, CNG channels, TRP channels, GPCRs, and beta-tubulins. CCSs overlapping target loci were extracted with BEDTools [33] and manually assessed with IGV [50] and R.

## Results

### Summary of sequencing results

We extracted high-quality RNA from *B. malayi* and *D. immitis* adult worms and subjected it to long-read sequencing with PacBio Iso-Seq. A single *D. immitis* worm generated sufficient quantities of RNA, while multiple *B. malayi* worms were pooled to achieve adequate RNA input. Sequencing of the four libraries and data preparation with IsoSeq3 resulted in a combined 160 Mb of high-quality isoforms from almost 134,000 CCSs. More detailed sequencing statistics can be found in S1 Table.

We assessed the quality of CCSs with SQANTI [30], which reference-corrects, filters, and categorizes putative isoforms based on a number of quality control metrics, and we removed CCSs that had characteristics of reverse transcriptase template switching. After filtration, 60,230 and 71,267 CCSs remained from *B. malayi* and *D. immitis*, respectively (Fig 1A). We identified CCSs mapping to 1,847/1,873 known genes in females, 1,651/1,433 in males, and 2,430/2,423 in both sexes (Fig 1B), covering 54.2% and 44.6% of the *B. malayi* and *D. immitis* coding genomes, respectively. Of these, we identified CCSs mapping to 6,197/10,522 unique transcripts in females, 6,646/5,542 in males, and 2,958/2,833 in both sexes (Fig 1C). These tallies of unique transcripts are partially inflated by the inclusion of truncated transcripts caused by RNA degradation, as we detected a 3’ bias in libraries from both species (Fig 1D).

**Fig 1.**
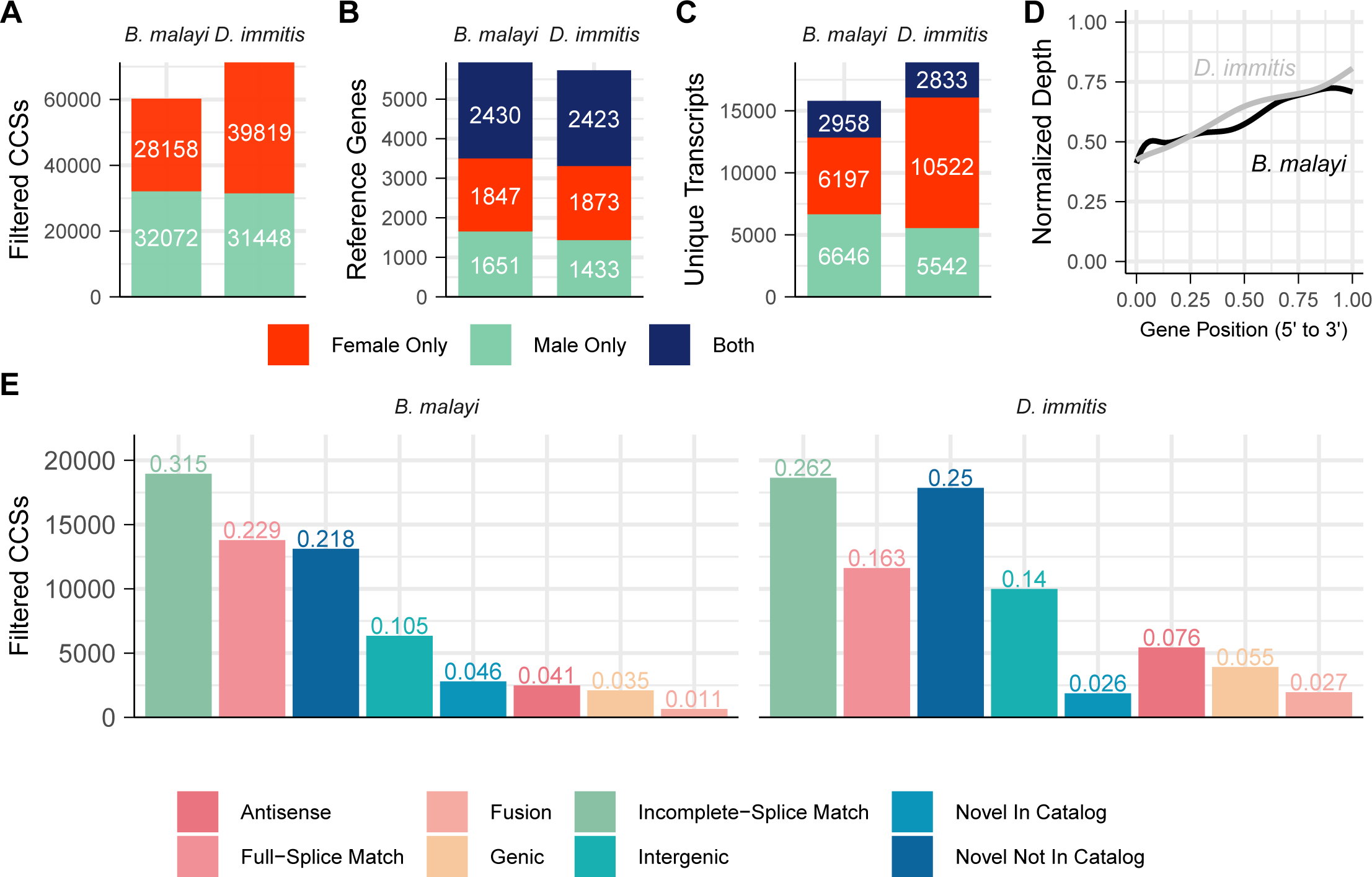
Summary of Iso-Seq data from adult male and female *B. malayi* and *D. immitis*. (A) Number of filtered CCSs generated from each sex and species. (B) Number of reference genes with at least one mapped CCS from only females, only males, or both sexes. (C) Number of unique transcripts captured from only females, only males, or both sexes. (D) 3’ bias in CCSs from both *B. malayi* and *D. immitis* due to poly(A) selection and mRNA degradation. (E) Distribution of SQANTI structural category assignments. Antisense: maps to the opposite strand of a predicted gene. Full-splice match: confirms the annotated gene model. Fusion: maps to two consecutive genes. Genic: maps to a single exon within a predicted gene. Incomplete-splice match: partial confirmation of the annotated gene model. Intergenic: maps to an intergenic region. Novel in catalog: utilizes known splice sites in novel combinations. Novel not in catalog: uses novel splice sites.

### Improvement and correction of gene models by Iso-Seq

We used SQANTI categories to identify gene models that were improved, corrected, and/or validated by our dataset (Fig 1E). Full-splice matches (FSMs) indicate models that were validated, and many of these CCSs extended UTRs. 57/77% extended the 5’ UTRs in *B. malayi* and *D. immitis*, respectively (Fig 2A), while 68/89% extended the 3’ UTR (Fig 2B). CCSs categorized as incomplete-splice matches (ISMs) were primarily 3’ fragments, but these were still able to extend gene models at the 3’ end (59/75% in *B. malayi* and *D. immitis*, respectively) (Fig 2B). In total, 1,632/3,713 transcripts in *B. malayi* and 1,481/3,502 in *D. immitis* had 5’/3’ UTRs extended.

**Fig 2.**
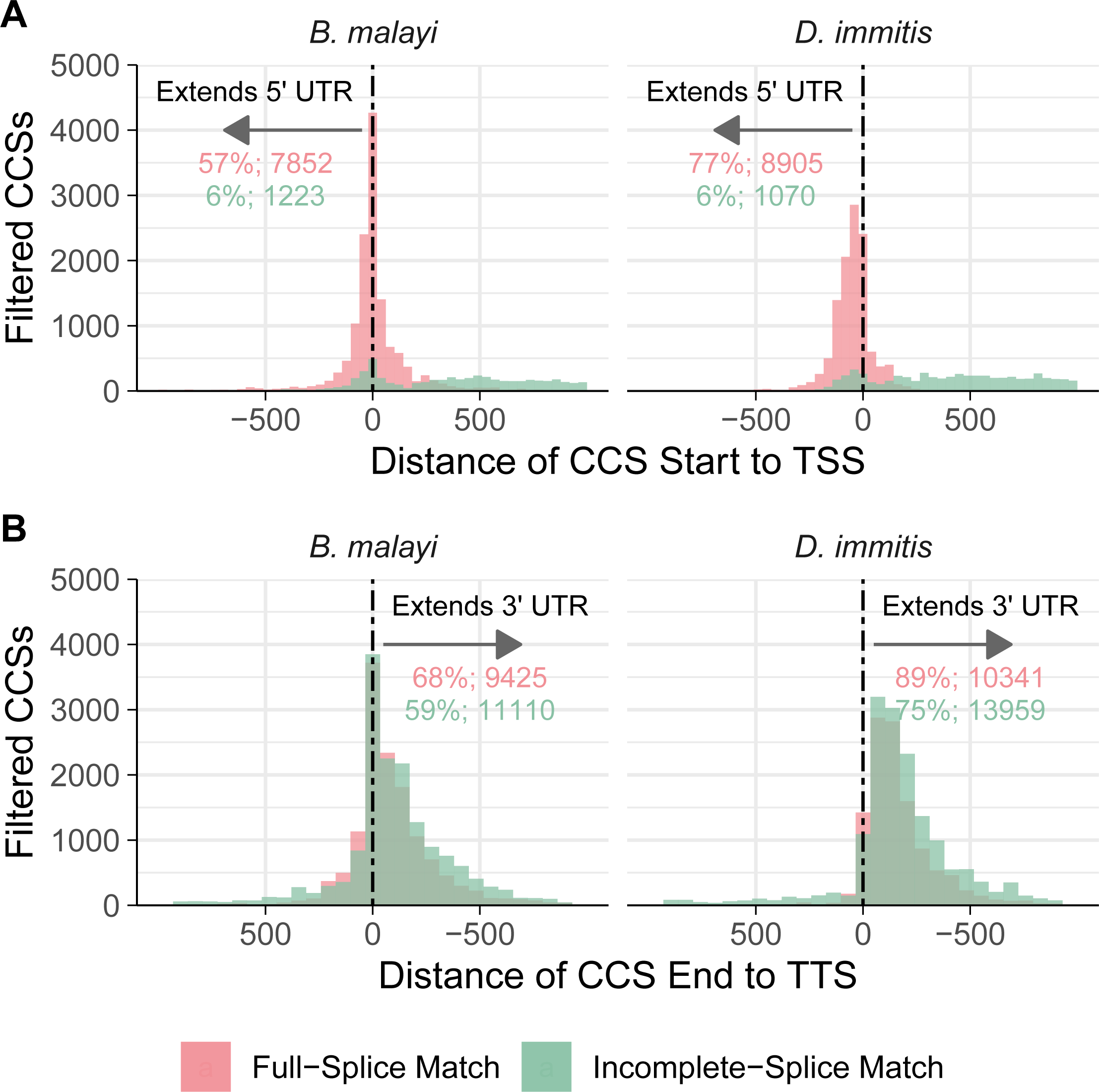
Iso-Seq extends UTRs of known transcripts. Filtered CCSs categorized as full or incomplete-splice matches were examined for whether they extended the 5’ and 3’ untranslated regions. (A) A high portion of full-splice match (FSM) CCSs extend the annotated 5’ UTRs of known transcripts in both *B. malayi* (left) and *D. immitis* (right), while incomplete-splice match (ISM) CCSs rarely cover the 5’ end. (B) Both FSM and ISM CCSs extend the 3’ UTRs of known transcripts. Negative values indicate CCSs that cover the transcriptional start or termination sites.

In addition to extended UTRs, we also added to the catalog of splice junctions used by these two filarial species. We identified 38,973/33,904 known and 4,527/4,651 novel canonical (G[UC]-AG) splice junctions and a total of 33/7 known and 818/947 novel non-canonical splice junctions in *B. malayi* and *D. immitis*, respectively (the top 10 splice junctions from *B. malayi* are shown in Fig 3A).

**Fig 3.**
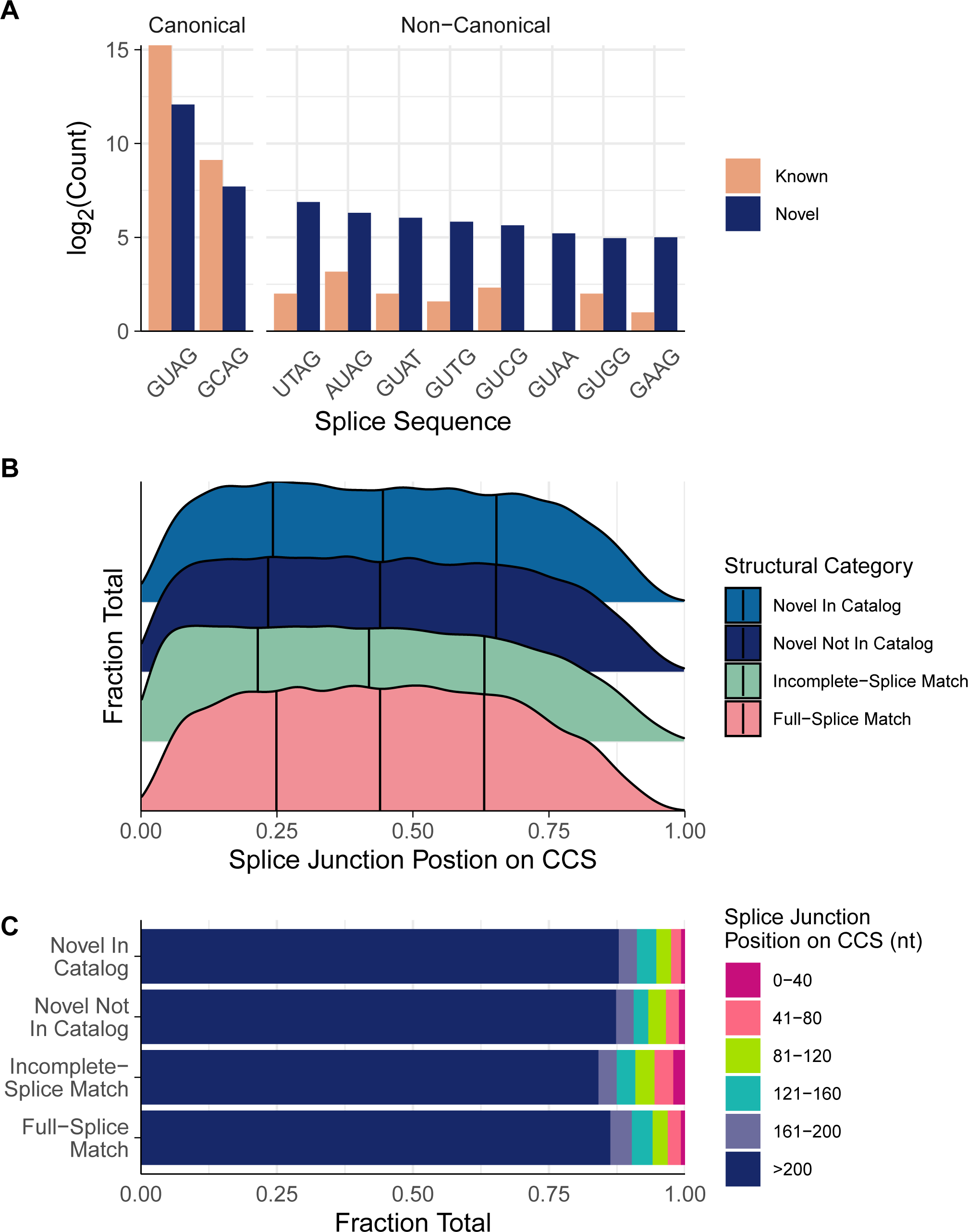
Iso-Seq adds to the catalog of splice junctions and identifies unannotated isoforms. (A) Most splice junctions mapped by CCSs were either the canonical G[UC]-AG, but a large portion of novel splice junctions identified were non-canonical. (B and C) Incomplete-splice match (ISM) CCSs contain a higher proportion of splice junctions at the 5’ end of the CCS when compared to full-splice matches (FSM) CCSs, novel in catalog (NIC) CCSs, and novel not in catalog (NNC) CCSs. This suggests that ISMs represent degraded mRNA, while NIC and NNC CCSs likely represent true novel isoforms or transcripts with corrected splice sites.

Finally, we identified CCSs that correspond to gene models that require concatenation (i.e., represent individual genes but are fragmented into multiple models). We identified CCSs that aligned to >1 gene, extracted sequences from the aligned genes, performed a search of the predicted proteins against the *C. elegans* predicted proteome, and looked for CCSs that had a single hit spanning the majority of the CCS (see S1 Fig for an example). These data were used to distinguish incorrectly fragmented gene models and putative operons, discussed in subsequent sections.

### Discovery of novel genes encoding proteins and potential long non-coding RNAs

In addition to identifying ways in which currently annotated gene models were improved upon, we used these data to search for entirely novel genes and isoforms. CCSs categorized as NIC or NNC align to either new combinations of splice junctions or novel unannotated junctions, making these categories rich in potential new isoforms. In known genes with high-quality models, splice junctions are equally abundant across the middle portion of transcripts (as shown by the FSM data in Fig 3B). Interestingly, junctions covered by NNC and NIC CCSs show a similar distribution to those covered by FSMs, while junctions covered by ISM CCSs are more abundant within the first 40 nt, causing a statistically significant 1.5% shift in mean position (p < 10^−10^, Tukey’s honest significant difference). CCSs classified as ISMs tend to be 3’ fragments, explaining why splice junctions had a greater propensity to map to these first 40 nt (Fig 3C). In contrast, the similar splice junction distributions among FSMs, NNCs, and NICs suggest that many NNC and NIC CCSs are full-length transcripts that represent unannotated isoforms.

We next looked for CCSs that may represent novel genes. We found a high number of CCSs categorized as antisense or intergenic, particularly in *D. immitis* (Fig. 1E). These categories represent CCSs that map to regions of the genome that do not have predicted genes (intergenic) or have predicted genes on the opposite strand (antisense). We found 1670/1878 multi-exonic loci of these potentially novel genes in *B. malayi* and *D. immitis*, respectively. We reasoned that these CCSs could represent either novel protein-coding genes or genes for non-coding RNAs (ncRNAs).

We focused on potentially novel protein-coding genes by filtering for CCSs that had a multi-exonic open reading frame (ORF), as mono-exonic CCSs are more likely to be the result of oligo(dT) priming of poly(A) stretches within contaminating genomic DNA. We found 209/501 of these CCSs mapping to 50/138 loci in *B. malayi* and *D. immitis*, respectively (S2 Table), and these CCSs are not similar to already predicted proteins but are similar to predicted proteins from other nematodes. Some of these putative genes were found to be specific to clade III nematodes, which may explain why they had not been predicted in *B. malayi*, given the heavy reliance of gene prediction algorithms on evidence from other nematode clades.

The remaining antisense and intergenic CCSs could represent ncRNAs. Only 39 ncRNAs are annotated in the *B. malayi* genome, and none have been identified in *D. immitis*. For *B. malayi*, all of these are mono-exonic, even though some ncRNAs such as long non-coding (lncRNAs) can be multi-exonic and use the same splicing machinery as pre-mRNAs. We found 67 CCSs mapping to 4 of these annotated ncRNA genes (WBGene00221733, WBGene00255379, WBGene00230800, and WBGene00220274). We attempted to determine whether the remaining CCSs could represent lncRNAs, but lncRNAs are typically expressed in lower abundance than mRNAs, and predicting them would require the integration of a large amount of short-read RNA-seq datasets combined with *de novo* gene prediction.

### Identification of new operons

A mixture of experimental and computational approaches have been used to annotate operons in the *B. malayi* genome, relying heavily on evidence from *C. elegans* [51,52]. Iso-Seq captures transcripts at a variety of processing steps, enabling the capture of some unspliced polycistronic transcripts. For operon prediction, we focused on the *B. malayi* data, which was less likely to include CCSs that mapped to single genes with multiple, fragmented gene models. We identified 1,775 CCSs that aligned to 443 loci containing multiple unique, consecutive genes.

Manual inspection of these putative operons revealed that many were likely to be indicative of fragmented gene models or were false positives resulting from misalignment, CCSs aligning to the opposite strand of neighboring genes, or genomic DNA contamination. We manually curated all putative operons that had a maximum intergenic distance of 5000, 338 loci in total.

As expected, we found that many predicted operons, both in our dataset and in the assembly annotations, were actually fragmented models that belonged to a single gene. These fragments tended to include one long fragment that was highly similar, but shorter, than its nearest nematode ortholog, and a second shorter fragment with an ORF of 30-60 amino acids. CCSs spanning these fragments had a single ORF that was significantly similar to a single nematode ortholog. We found 89 of these fragmented loci that require concatenation, and 31 of these are annotated operons in the Bmal-4.0 assembly, which should be deprecated. On the other hand, true operons had ORFs that were of similar size and typically >100 amino acids, and the CCSs spanning them often had unspliced intercistronic introns that separated two large ORFs with similarity to multiple nematode orthologs. We identified 31 novel operons and validated 102 assembly-annotated operons. These 133 operons contain 382 genes for an average of 2.87 genes per operon (Fig 4A, S3 Table). We pulled *C. elegans* orthologues from WormBase ParaSite [28] and found that most of the newly identified operonic genes do not have a *C. elegans* orthologue (Fig 4B). This is likely explained by the previous utilization of *C. elegans* operons to inform *B. malayi* operon annotation, highlighting the power of unbiased sequencing approaches to identifying operons. The distances between consecutive cistrons in *B. malayi* operons are substantially shorter than monocistronic genes (Fig 4C, 1766.31 nucleotides for cistrons and 16821.48 for non-operonic genes), as in other nematodes, but the distribution of distances is much broader than that of *C. elegans* (Fig 4D, interquartile range of 1363 contrasted to 700.5 for *C. elegans*). This may be explained by the lack of the SL2 splice-leader sequence in *B. malayi*. While SL2 is the predominant SL used for trans-splicing of downstream cistrons in *C. elegans*, SL1-spliced downstream cistrons are often associated with longer intergenic distances [53].

**Fig 4.**
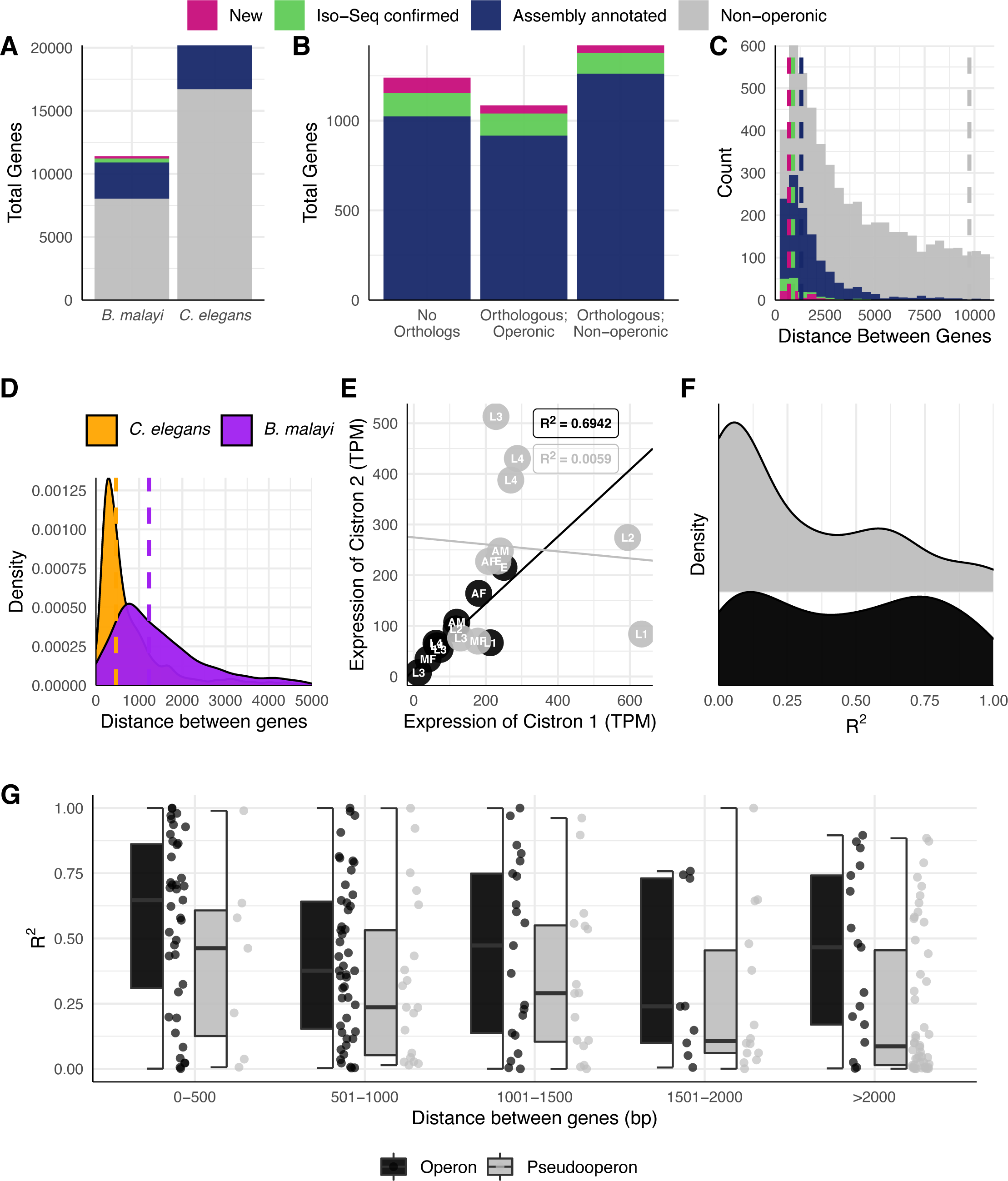
Identification and validation of over a hundred operons in *B. malayi*. (A) Iso-Seq identified new operons and validated assembly-annotated operons in *B. malayi*, which has a similar proportion of its genome arranged in operons as *C. elegans*. (B) Most genes in novel operons do not have orthologs in *C. elegans*. (C) Intercistronic distances between genes in operons are much shorter than intergenic distances between non-operonic genes. Operons identified via Iso-Seq (974.09 nucleotides for new operons, 1197.01 for validated operons) on average had shorter intercistronic distances than those already annotated in the Bmal-4.0 assembly (1879.23). (D) *B. malayi* operons on average have larger (median = 1227.5 contrasted to 467 for *C. elegans*) and a broader range (interquartile range of 1363 contrasted to 700.5 for *C. elegans*) of intercistronic distances than those in *C. elegans*. (E) An example of linear fits to staged RNA-seq data from genes in a pseudo-operon (non-operonic neighboring genes) and an operon. (F) Linear fits better explain the relationship between expression of neighboring genes in operons than neighboring genes in pseudo-operons, and (G) this contrast does not stratify by intergenic distance.

Operons allow for the co-regulation of neighboring genes, which often results in transcript abundances that are correlated between paired cistrons [54]. We found that the expression abundances of the first pair of neighboring cistrons show a much stronger linear relationship than pseudo-operons created by neighboring monocistronic genes (Fig 4E-G). We used publicly available short-read RNA-seq data from staged parasites and fit a linear model to these values using the abundance of the first cistron as a predictor for the abundance of the second cistron (example shown in Fig 4E). In general, a linear model did not fit well to consecutive monocistronic genes (median R^2^ = 0.14), but performed much better for operonic genes (median R^2^ = 0.475) (Fig 4F), and this contrast is not altered by the intergenic distance (Fig 4G). Interestingly, the R^2^ values of the linear fits of operonic genes appear to be bimodal, which could indicate post-transcriptional regulation that may cause a deviation from the linear relationship.

### Analysis of poly(A) tails in filarial nematodes identifies alternative poly(A) addition sites

PacBio chemistry is adept at sequencing long homopolymers, allowing for the analysis of poly(A) tails and poly(A) addition sites (PAS). Though there are alternative focused methods for determining poly(A) tail lengths, derivatives of Iso-Seq have been shown to correlate well in *C. elegans* [55]. We found the median poly(A) tail length for CCSs from *B. malayi* was 33 nt (range of 1-541) and 35 (range of 1-536) for *D. immitis* (Fig. 5A), shorter than the reported *C. elegans* median of around ∼50 nt (depending on life stage and technique) and much shorter than the typical lengths for human organoids, iPSCs, and cell lines [55–57]. Nearly 40% of *B. malayi* CCSs contained the canonical AAUAAA PAS motif, with AUAAAN as the second most abundant (approximately 15% in total) (Fig. 5B). About 8% of CCSs did not have an identifiable PAS but instead included an A-rich region in the 50 nt upstream of the poly(A) addition site (Fig. 5B-D). These patterns were shared in *D. immitis* PASs (S2-5 Figs). Interestingly, we also identified an upstream sequence element (USE) that is a known recognition site for some RNA binding proteins (UUUUGU) in approximately 4% of CCSs that did not have other PASs [58,59].

**Fig 5.**
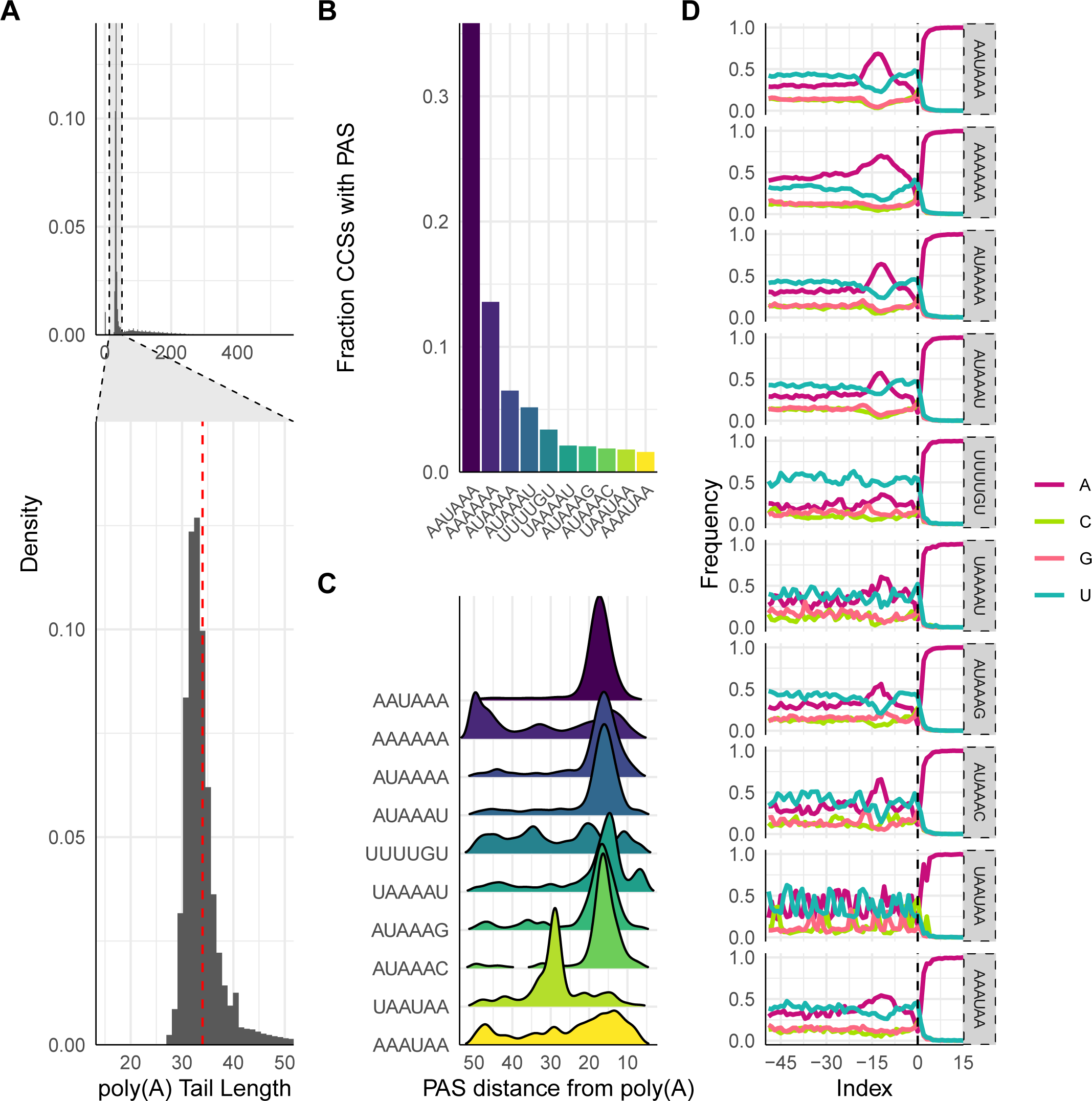
Analysis of poly(A) tails in *B. malayi*. (A) Poly(A) tails on CCSs from *B. malayi* ranged from 0 - 600 and had a median length of 33. (B) Greater than 35% of CCS contained the canonical poly(A) signal (PAS) AAUAA, while A-rich regions were the second most abundant PAS. (C) All PASs occur 10-20 nucleotides upstream of the poly(A) cleavage site, other than AAAAA, which has two A-rich sites at 10-20 and 45-50 nucleotides upstream, and UAAUAA, which may not be a PAS but is a motif for RNA binding proteins. (D) All PASs also include an A-rich region at 15 nucleotides upstream of the cleavage site.

### Improvement of gene models for known and putative anthelmintic targets

We curated a list of 97 genes of interest in *B. malayi* and *D. immitis*, consisting of known anthelmintic targets and receptors belonging to druggable protein families in parasitic nematodes (S4 Table). We examined the genomic loci of these targets for CCSs that mapped to these locations and cross-referenced these CCSs with SQANTI classifications. In *B. malayi*, 28 of these targets had mapped CCSs, most with multiple CCSs, and 11 of these had FSMs that confirmed the reference model and included 5’ and 3’ UTRs. For example, *Bma-slo-1* (Bm6719) had FSMs from both sexes (Fig. 6A) and *Bma-gar-3* (Bm13584) had the b isoform validated (Fig. 6C). *Bma-slo-1* has been shown to have sex-specific splice patterns that likely contribute to the differential response to emodepside [21], and our Iso-Seq data detected the usage of both the a and f isoforms in females and only the f isoform in males, consistent with this interpretation (Fig. 6A). We also captured in females an additional isoform that contains both the exons that show alternative usage in the a and f isoforms; this isoform would not have been picked up by PCR and restriction digest test used to previously define *Bma-slo-1* isoform usage.

**Fig 6.**
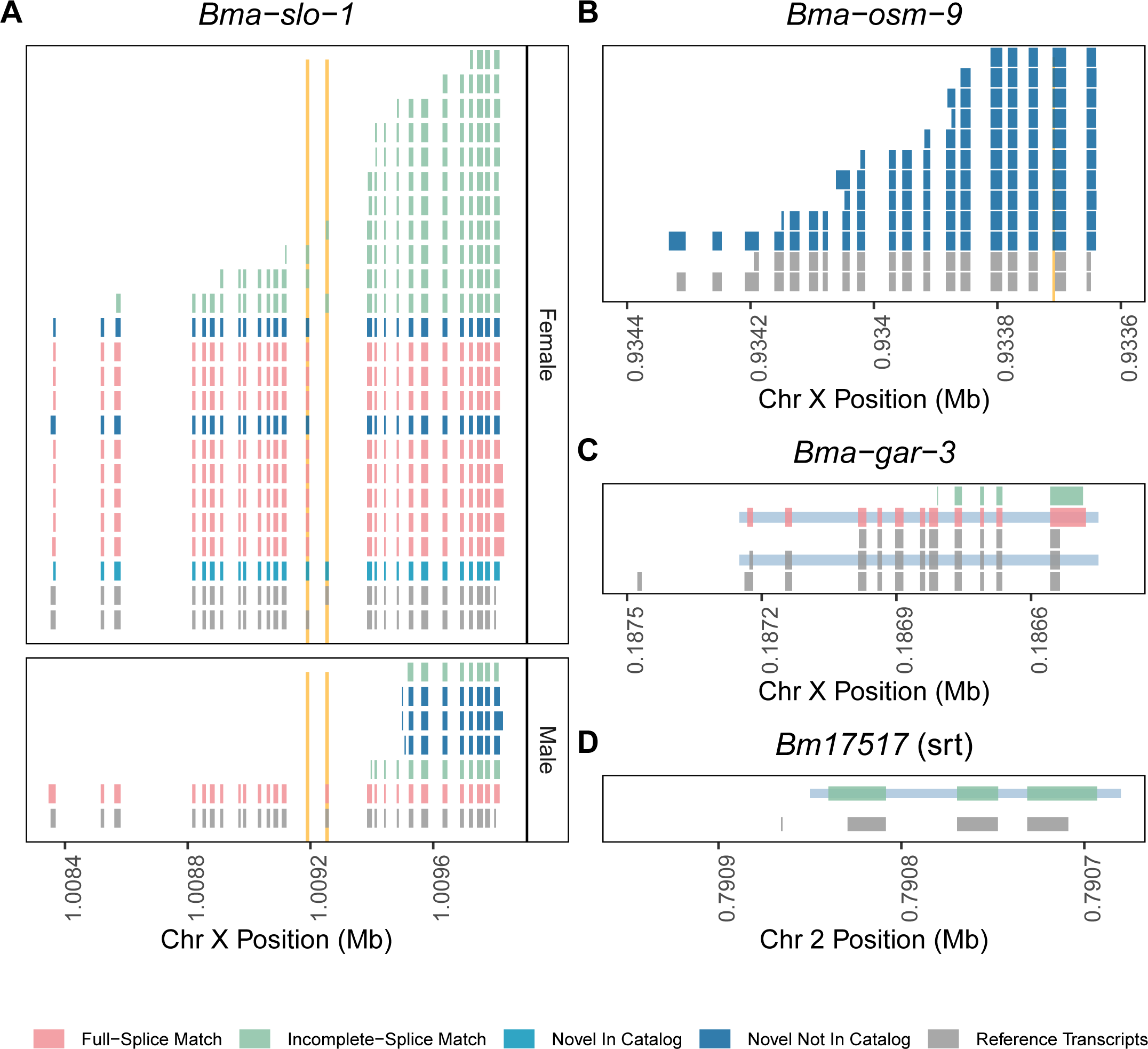
Correction of models for genes currently studied as potential anthelmintic targets. (A) *slo-1*, a target for emodepside, has multiple alternative isoforms that play a role in emodepside response [21]. Iso-Seq captured a previously unidentified isoform in *B. malayi* females. (B) All CCSs mapping to *osm-9* showed a mispredicted splice junction in the penultimate exon. (C) Iso-Seq confirms the beta isoform of *gar-3*. (D) Iso-Seq shows the incorrect prediction of the first exon and transcriptional start site of *Bm17517*, a chemoreceptor, and extends the coding sequence of exon 1.

Other targets had models corrected by mapping CCSs. While only a single full-length *Bma-osm-9* CCS was captured, multiple CCSs from degraded transcripts validated the previously corrected splice recognition site in *Bma-osm-9* (Fig. 6B) [60]. Finally, a read mapping to *Bm17517* (an *srt* family chemoreceptor [60]), showed evidence for a corrected or alternative TSS and start codon (Fig. 6D). The reference model for *Bm17517* contains a 9 nt first exon that has limited evidence from short-read and EST data sources, while Iso-Seq CCS suggests that this exon is spurious and instead the ORF of the second exon extends further upstream. These data demonstrate the ways in which Iso-Seq data can be used to correct the models for genes that are under active research and cloning efforts. The remaining 24 targets with mapped CCSs (including acetylcholine and glutamate-gated ion channels, TRP channels, CNG channels, and GPCRs) can be found in S2-5 Figs.

## Discussion

In recent years there has been a concerted push to increase the availability of genomic resources for parasitic worms of medical, agricultural, and veterinary importance, highlighted by the recent release of 81 draft genomes from both free-living and parasitic flatworms and nematodes [35]. These resources promise to considerably bolster our understanding of the evolution of parasitism and parasitic biological processes through the use of comparative genomics.

Although many important analyses can be performed with draft genomes of suboptimal contiguity, experimentation upon individual genes requires gene model predictions in which one can be highly confident. *Ab initio* gene prediction algorithms can be optimized to provide high BUSCO or CEGMA scores but often miss the genes in which parasitologists are most interested: those that are not conserved in hosts and are likely to be involved in parasitism [61,62].

Thus, the next substantial push for curators of helminth genomes should be to improve the gene model prediction and annotation pipeline. Short-read RNA-seq data has been instrumental in this aim and will continue to play an important role, but the technology is limited in its ability to differentiate between isoforms or resolve full-length transcripts. The recent development of long-read sequencing technologies can overcome these obstacles.

We report here the first long-read RNA-seq data for filarial nematodes, an important group of parasites that infect humans and animals. While the genome assembly of *B. malayi* is chromosome-scale, there remains serious errors in gene models, even for genes that are highly conserved among nematodes and have been cloned in other species [60]. *D. immitis*, on the other hand, has a far less contiguous genome assembly likely contains even more mispredicted gene models than *B. malayi*. The completeness of the *B. malayi* and *D. immitis* transcriptomes stands to be significantly improved once these data are integrated into gene and transcript prediction pipelines. The transcriptome assembly of *D. immitis*, in particular, will be substantially improved by these data. Until now, there was not a single alternative isoform predicted in this species, predicted UTRs were sparse, and very few non-canonical splice sites were annotated. Filtered CCSs for *B. malayi* have been included in WormBase (WS277) as genome browser tracks, and these data will be integrated into gene prediction pipelines in future releases.

Our data had a high 3’ bias due to RNA degradation and poly(A) selection and covered approximately 50% of the coding transcriptomes of *B. malayi* and *D. immitis*. Recent improvements in the Iso-Seq library preparation workflow will increase transcriptome coverage and isoform discovery, and allow for the sequencing of RNA from RNA-poor larvae and individual worms or tissues. A reduction in minimum RNA input will also enable techniques that enrich for full-length RNA molecules that have both the m7G cap and the poly(A) tail [63,64]. These approaches promise to further improve upon the enhancements we report here and should be used in the cases of other nematode species.

## Acknowledgements

Some parasite materials were provided by the NIH/NIAID Filariasis Research Reagent Resource Center (www.filariasiscenter.org). *D. immitis* RNA extractions were carried out by Jessica Grant and Steve Williams at Smith College. RNA sequencing was carried out at the University of Wisconsin-Madison Biotechnology Center. Data was uploaded to WormBase by Michael Paulini (WormBase). The authors would like to thank members of the Zamanian laboratory for critical comments on the manuscript.

## Funding

Funding for MZ is provided by an R01 (R01AI151171, NIH.gov) grant. The funders had no role in study design, data collection and analysis, decision to publish, or preparation of the manuscript.

## Supporting Information

**S1 Fig. Example of a fragmented gene model (*Dim-tax-4*) that requires concatenation**.

**S2 Fig. CCSs mapping to the two *ben-1* orthologs in *B. malayi***.

**S3 Fig. CCSs mapping to glutamate and acetylcholine-gated ion channels in *B. malayi***.

**S4 Fig. CCSs mapping to miscellaneous ion channels in *B. malayi***.

**S5 Fig. CCSs mapping to GPCRs in *B. malayi***.

**S1 Table. Sequencing statistics and metadata**.

**S2 Table. List of genomic loci in *B. malayi* and *D. immitis* that may contain novel genes**.

**S3 Table. List of validated and novel operons and their genes in *B. malayi***.

## Notes

### Competing Interest Statement

The authors have declared no competing interest.

