## Supplementary figures and images for "Long-read RNA sequencing of human and animal filarial parasites improves gene models and discovers operons"

### S1 Fig

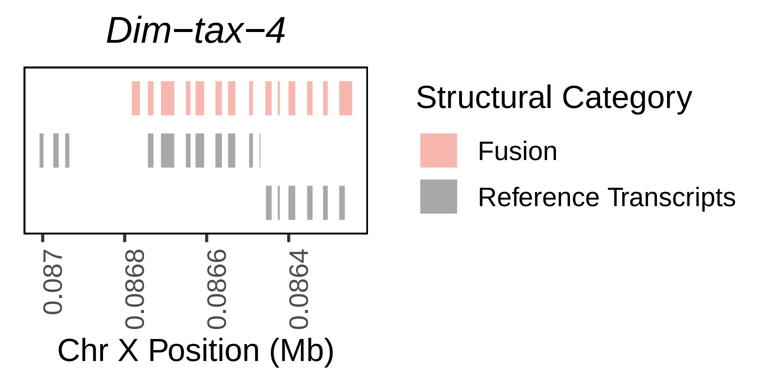

### S2 Fig

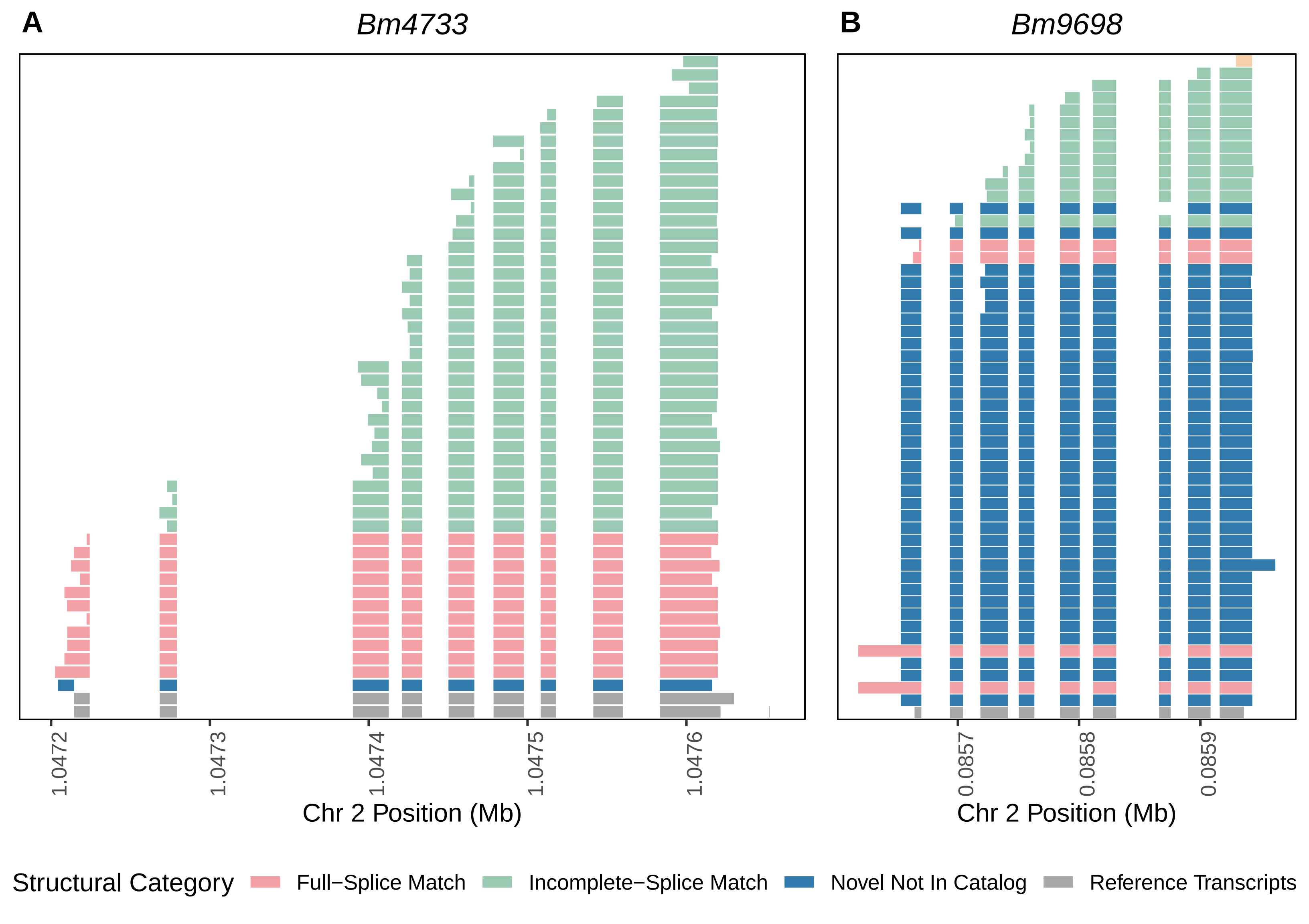

### S3 Fig

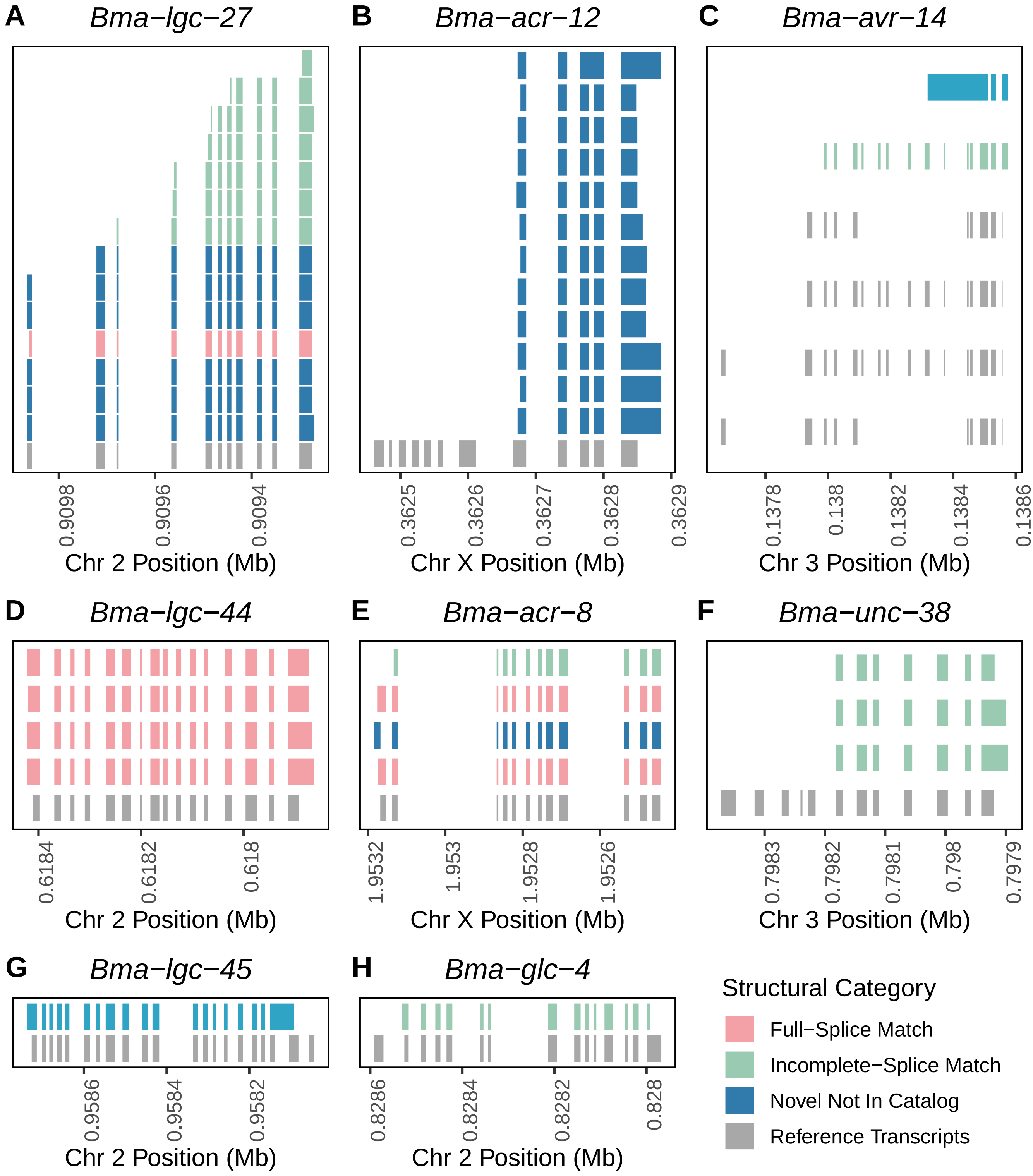

### S4 Fig

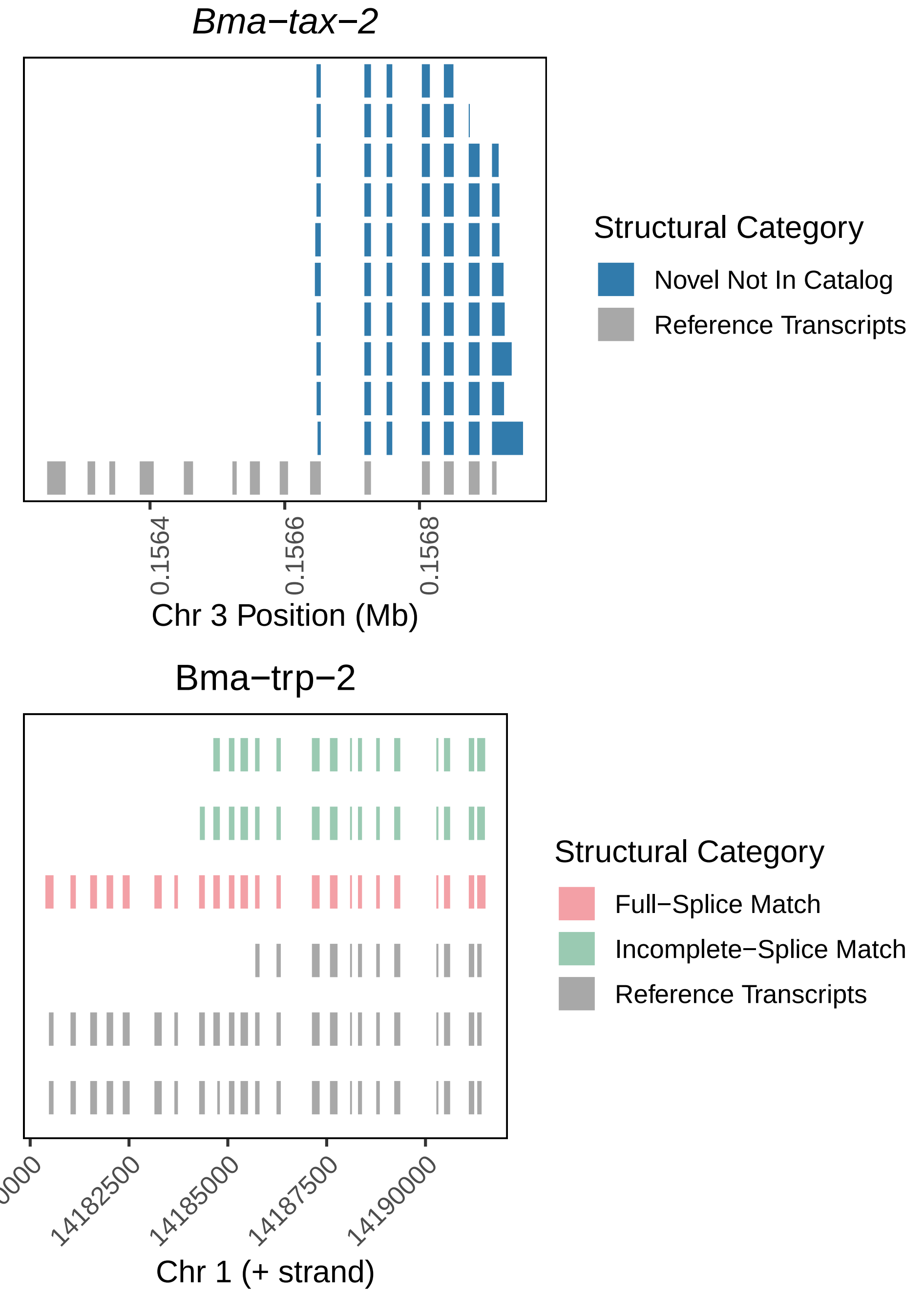

### S5 Fig

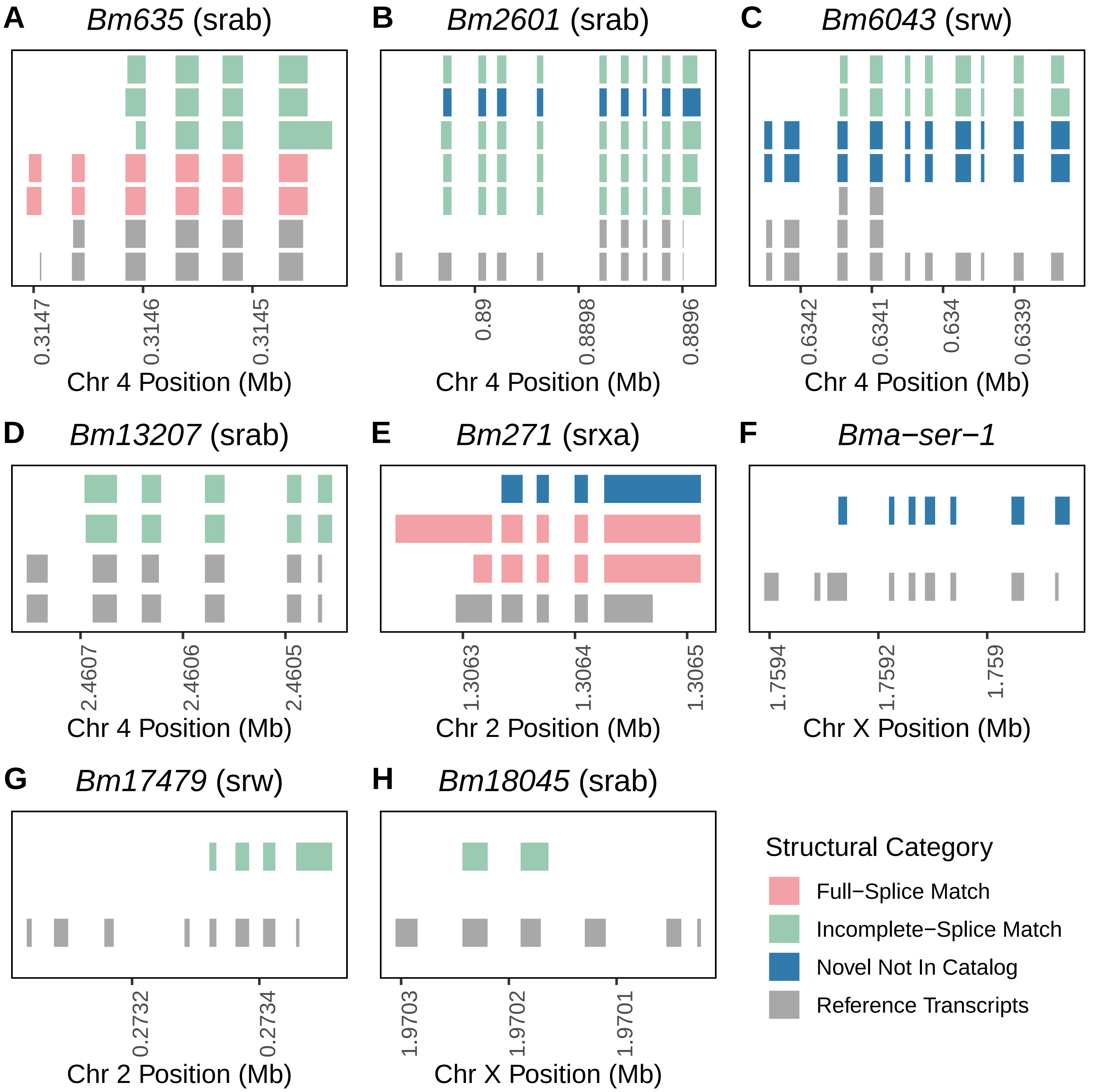
